# A comparative, multi-study analysis of plastisphere resistomes, plasmid dynamics, and antibiotic resistance genes

**DOI:** 10.64898/2026.01.11.698908

**Authors:** Isaac J. Okyere, Chanistha Tiyapun, Jennifer L. Goff

**Author notes:** Corresponding Author; 1 Forestry Dr., Syracuse, NY 13210.

## Abstract

Microplastics are widespread in aquatic environments and support surface-associated microbial communities. Although antimicrobial resistance in the plastisphere has been reported, the organization of resistance genes across plasmid types and mobility categories on microplastic surfaces remains incompletely characterized. In this study, we performed a comparative analysis of published microplastic biofilm metagenomes to examine plasmid replicon diversity, predicted mobility, and antimicrobial resistance gene (ARG) distributions across various microplastic polymers. Phylum-level taxonomic profiles varied across polymer types, but most plastisphere communities were dominated by *Pseudomonadota*. Plasmid reconstruction revealed differences in predicted mobility profiles, with conjugative plasmids more frequently associated with polyethylene, polypropylene, and polyvinyl chloride. Network analyses linking plasmid mobility categories, replicon types, and ARG classes showed that conjugative plasmids connected a broader range of replicons to multiple ARG classes than mobilizable or non-mobilizable plasmids. Replicons from the Inc family, especially IncFIB and IncFII, were prominent and carried high ARG loads, consistent with their capacity to harbor multiple accessory genes. Additionally, several less-characterized rep_cluster replicons were detected across microplastic types, indicating diverse and understudied plasmid backbones within plastisphere communities. The co-occurrence of ARGs with mobile plasmid architectures underscores the importance of considering plasmid context when evaluating plastisphere resistomes.

**IMPACT STATEMENT:** Microplastics are increasingly recognized as microbial habitats that can concentrate antibiotic resistance genes (ARGs) in aquatic environments. While many studies have documented the presence of ARGs within microplastic-associated biofilms, far less is known about the genomic context of these genes. This study advances the field by shifting the focus from simple ARG inventories to the plasmid architectures associated with these ARGs. By integrating metagenomic data from previously published studies spanning freshwater, estuarine, and marine systems, we provide the first comparative, cross-system assessment of plasmid replicon diversity, mobility potential, and ARG co-occurrence across different microplastic polymers. This plasmid-centric perspective reveals that ARGs in microplastic biofilms are often associated with conjugative plasmids, which can facilitate their horizontal transfer within these biofilms. Importantly, this work identifies microplastics as environments where clinically relevant ARGs are linked to plasmid architectures commonly observed in pathogenic bacteria. This conceptual advance supports more mechanistic risk assessments of plastic pollution and informs One Health-oriented strategies to address antibiotic resistance across environmental, animal, and human systems.

## INTRODUCTION

Antibiotic resistance has emerged as a global health crisis, driven mainly by the extensive use of antibiotics in human medicine and livestock farming (Aslam *et al*. 2021; Burow and Käsbohrer 2017; Chaw *et al*. 2018). Compounding this challenge, no new classes of antibiotics have been discovered in decades, leaving humans and animals vulnerable to infections that were once treatable (Durand *et al*. 2019; Miethke *et al*. 2021). It is currently estimated that antibiotic resistance is responsible for approximately 700,000 deaths annually, and this number is projected to rise to 10 million deaths per year by 2050 if no effective interventions are implemented (Tagliabue and Rappuoli 2018). While much attention has been given to clinical and animal sources of antibiotic resistance, growing evidence indicates that natural environments, such as aquatic systems, are significant contributors to the spread and persistence of antibiotic resistance. Aquatic ecosystems have become reservoirs of antibiotic-resistant bacteria (ARB) and their associated antibiotic resistance genes (ARGs), enriched by the influx of untreated or partially treated waste such as municipal wastewater, pharmaceutical effluents, and agricultural runoff (Ferreira da Silva *et al*. 2006; Ouyang *et al*. 2015). These contaminants introduce antibiotics and resistant microbes into water bodies, creating hotspots where resistance can evolve and spread (Baquero *et al*. 2008; Ma *et al*. 2015).

Alongside these chemical and microbial pollutants, plastic waste accumulation in aquatic environments raises additional concerns. Plastics, produced in large quantities for industrial and consumer use, are highly durable and resistant to degradation (Fayshal 2024). Over time, larger plastic debris breaks down into smaller fragments known as microplastics, which are defined as plastic particles less than 5 mm in size (Amaral-Zettler *et al*. 2015; Bajt 2021). These environmental contaminants have become ubiquitous in marine, freshwater, and terrestrial environments. Due to their small size, large surface area, chemical stability, and hydrophobicity, microplastics persist and often co-occur with ARBs and ARGs in aquatic environments, acting as substrates for microbial colonization (Gong *et al*. 2019; Reisser *et al*. 2014). This colonization leads to the formation of biofilm-associated microbial communities, often referred to as the “plastisphere” (Wu *et al*. 2019a; Zettler *et al*. 2013). The plastisphere is composed of diverse microbial communities, including opportunistic pathogens such as *Vibrio spp.*, *Aeromonas spp., Acinetobacter spp.,* and *Pseudomonas spp.*, all of which have been identified on microplastics in both marine and freshwater systems (Kirstein *et al*. 2016; McCormick *et al*. 2014; Oberbeckmann *et al*. 2016).

These colonized microplastics not only provide a habitat for microbial proliferation but also facilitate horizontal gene transfer (HGT), a process by which bacteria exchange ARGs, often mediated by plasmids, integrons, and transposons (Luo *et al*. 2023). Thus, the plastisphere can act as a hotspot for ARG exchange. Moreover, microplastics can travel long distances across water bodies, contributing to the dissemination of ARB and ARGs across ecosystems (Horton and Dixon 2018). These particles can then be ingested and bioaccumulated by aquatic organisms across various trophic levels (Miller *et al*. 2020). Through such pathways, they may ultimately reach humans via contaminated seafood or drinking water, a concern supported by evidence of microplastics detected in human feces (Yan *et al*. 2022). Thus, highlighting their emergence as a growing public health concern.

Metagenomic studies investigating plastisphere-associated microbial communities have revealed the complexity of resistome profiles. However, comparative, multi-study analyses remain limited, especially those that integrate plasmid dynamics, ARG profiles, and taxonomic diversity across different aquatic environments. To address this gap, we collected published metagenomic datasets generated from across different geographical locations and aquatic systems. These datasets were analyzed using a standardized bioinformatics pipeline to identify key microbial taxa within the plastisphere, assess ARG diversity, and characterize associated plasmid types.

## METHODS

### Data curation

In August 2024, we performed a literature search in Google Scholar using the key terms “microplastics”, “biofilm”, “metagenome”, and “aquatic” to identify metagenomic datasets generated from microplastic biofilms **(Table S1).** Specifically, we focused on metagenomes from microplastics collected from natural bodies of water (Bryant *et al*. 2016; Delacuvellerie *et al*. 2022; Di Cesare *et al*. 2024). Thus, we excluded engineered or built environment systems (*e.g.,* wastewater treatment plants). However, to expand the number of included studies, we did retain studies that set up bioreactor incubation experiments using microplastics with water from a natural system (Wu *et al*. 2022) or *in situ* incubations of microplastics in natural bodies of water (Bhagwat *et al*. 2021; Oberbeckmann *et al*. 2021). To ensure data relevance, each dataset was manually reviewed to confirm that the sequencing reads represented untargeted metagenomes and matched the metadata descriptions. Studies were excluded if the study metadata did not match the metadata in the sequence read archive (SRA), if the metagenomes were not generated from microplastic biofilms, or if the sequenced reads were not publicly available. Additionally, we excluded amplicon sequencing-based datasets. Only datasets meeting these criteria were retained for subsequent analyses.

### Bioinformatic analyses

The analyses described below were performed on the Department of Energy (DOE) KnowledgeBase (KBase) platform (Arkin *et al*. 2018) using the following pipeline with default parameters, unless otherwise specified. All publicly available raw sequence data from the selected studies were imported from the SRA using the Import SRA File as Reads from Web tool with the SRA identifiers compiled in our literature search. Initial quality assessments of the raw sequence reads were performed using FastQC (v0.12.1) (Andrew S. 2010) to detect low-quality bases and identify adapter sequences. Adapter sequences were first removed using Cutadapt (v4.4) (Martin 2011), after which reads were further trimmed, if needed, using Trimmomatic (v0.36) (Bolger *et al*. 2014) before the quality of the trimmed outputs was reassessed using FastQC (v0.12.1). We used metaSPAdes (v3.15.3) (Nurk *et al*. 2017) to assemble the trimmed reads to generate contig sets. These contigs were binned using MaxBin2 (v2.2.4) (Wu *et al*. 2014) to produce binned contig objects. In a mock community, metaSPAdes in combination with MaxBin2 was previously found to assemble and bin >60% of plasmids (with >50% coverage) (Maguire *et al*. 2020). As binning tools rely on GC content and coverage to link contigs together (and plasmids often diverge from their host genome in both these metrics), we relied on other tools to infer the host phylogeny of the plasmids recovered from bins, discussed later. Medium- and high-quality bins were then filtered using CheckM (v1.0.18) (Parks *et al*. 2015) with thresholds of ≥50% completeness and ≤ 10% contamination. Medium and high-quality bins were further processed to extract individual assemblies using the Extract Bins as Assemblies tool in KBase. The resulting assemblies were combined into a single file per study to streamline downstream analyses. Metagenomic reads from each sequencing library were mapped to the assembled MAGs using Bowtie2 (v2.3.2) (Langmead and Salzberg 2012), and the abundance of each MAG was calculated as the proportion of reads mapping to the MAG relative to the total number of reads in the corresponding library. Taxonomic classification of the MAGs was performed using GTDB-Tk (v2.3.2)(Chaumeil *et al*. 2019), which assigns taxonomy based on the Genome Taxonomy Database. Phylum-level relative abundances were obtained by summing the abundances of all MAGs assigned to each phylum and expressing the resulting values as percentages of the total microbial community.

These combined sequence files were then uploaded into the DOE National Microbiome Data Collaborative (NMDC)-developed Empowering the Development of Genomics Expertise (EDGE) platform (Kelliher *et al*. 2024) and analyzed using the Viruses and Plasmids workflow, which utilizes geNomad (v4.1.0) for mobile genetic element identification (Camargo *et al*. 2024). This workflow enabled the identification and characterization of plasmid-associated sequences within the MAGs. Plasmid-associated ARGs were identified using the Resistance Gene Identifier (RGI, v6.0.5) tool against the Comprehensive Antibiotic Resistance Database (CARD v4.0.1) (Alcock *et al*. 2023) with parameters set at selecting perfect, strict, and loose hits, and excluding nudge ≥95% identity loose hits to strict. Additional confirmation was done using AMRFinderPlus (v2024-01-31.1) (Feldgarden *et al*. 2021) Plasmid sequences obtained from geNomad were further screened against the Plasmid Database (PLSDB, v2024_) using Mash screen with parameters (max. p-value = 0.1, min. identity = 0.80) to predict known plasmids present in the input sequences (Molano *et al*. 2025). The summarized workflow used in this study is shown below in **Fig. 1**.

**Fig. 1.**
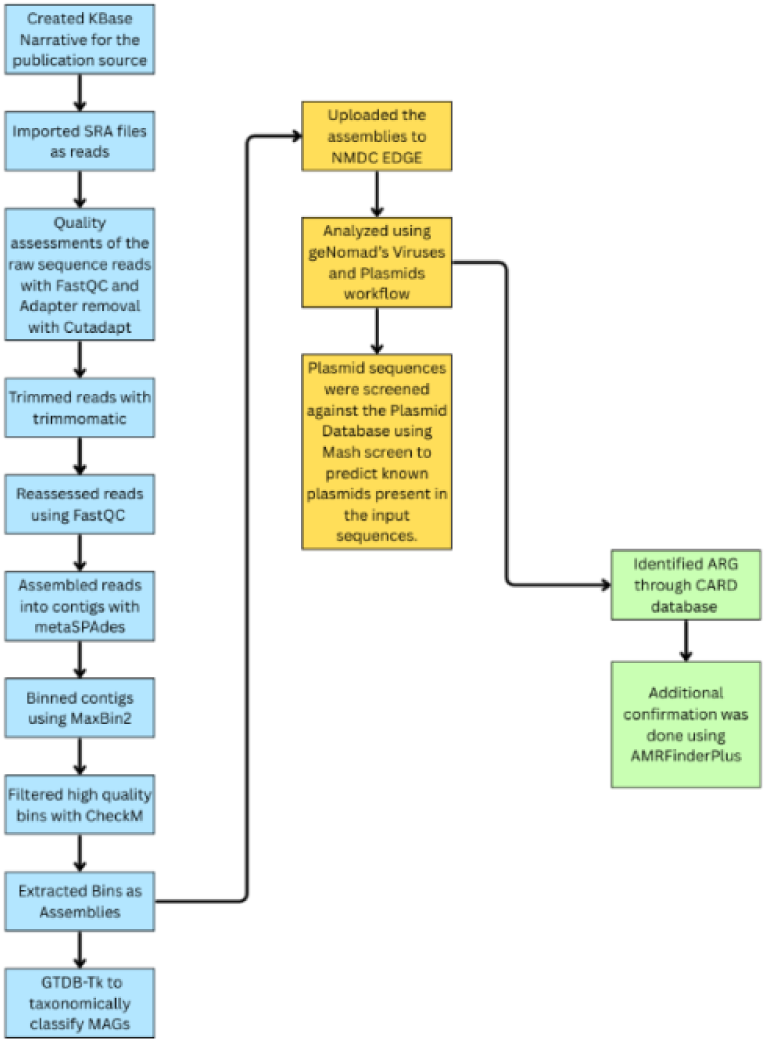
Workflow of metagenomic analysis for ARG and plasmid identification

## RESULTS

### Microplastics-Associated Bacterial Communities

We compiled metagenomic datasets from six independent studies investigating microplastic-associated microbial communities. Collectively, the datasets encompassed samples derived from both marine (n = 4 studies), estuary (n = 1 study), and freshwater (n = 1 study) environments. Geographically, the studies spanned three continents, with datasets originating from Europe (n = 3 studies: Baltic Sea/Warnow estuary, Tyrrhenian Sea, Mediterranean Sea), Asia (n = 1 study: urban river, China), Australia (n = 1 study: coastal marine, Australia), and the open ocean of the North Pacific Gyre (n = 1 study) (**Fig. 2**). In total, the combined dataset included 217 metagenomic read libraries. A total of 1,188 MAGs were binned. Of these, 341 were classified as medium quality (≥ 50% completeness and ≤ 10% contamination), while 105 met the criteria for high-quality MAGs (≥ 90% completeness and ≤ 5% contamination) **(Table S2).** Medium- and high-quality MAGs were retained for further analyses.

**Fig. 2.**
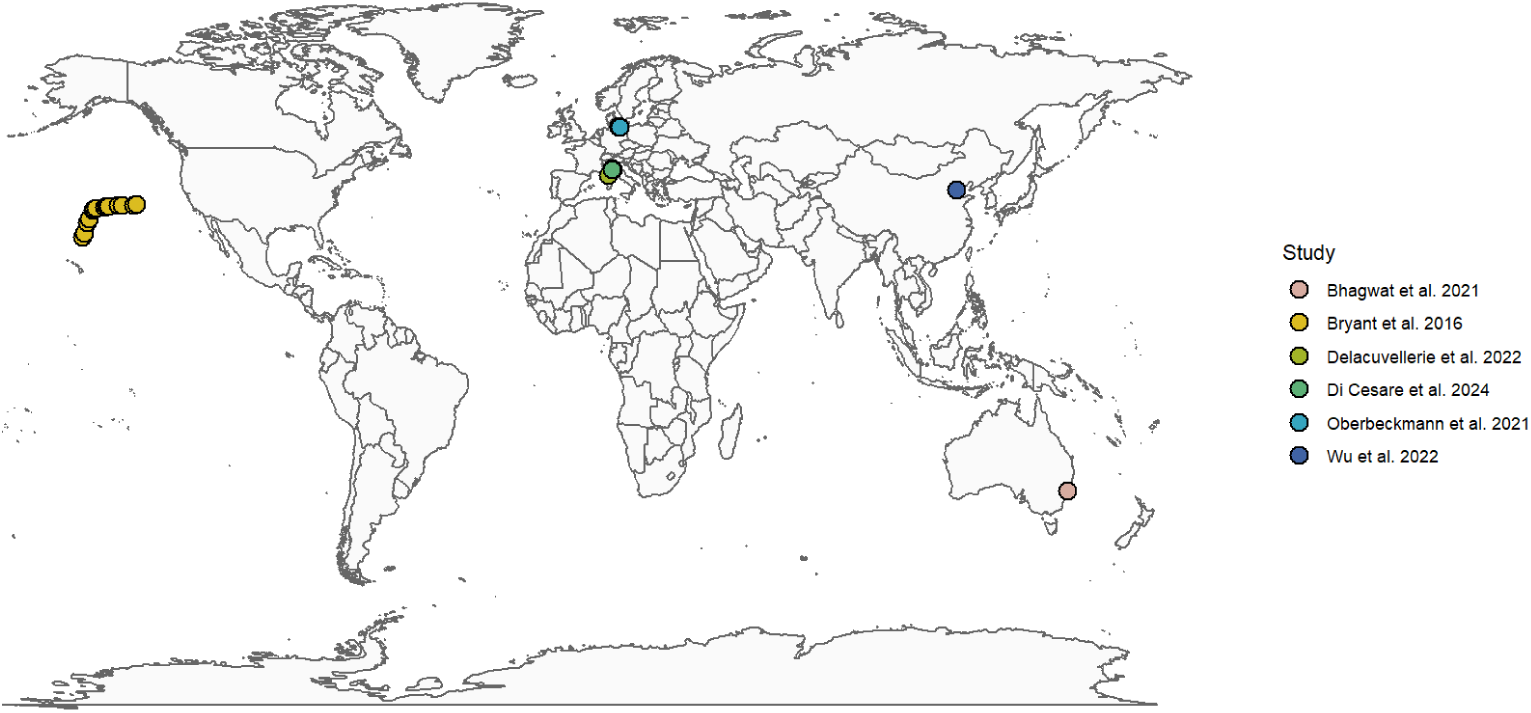
Map showing the geographical distribution of the metagenomic datasets included in this analysis. Sampling sites spanned Europe (Baltic Sea, Tyrrhenian Sea, Mediterranean Sea), Asia (Haihe River, China), Australia (Lake Macquarie estuary), and North America (North Pacific Gyre).

Polymer types associated with the microplastic biofilms were determined using metadata for the sequencing libraries retrieved from the SRA database. Each sequencing library was categorized according to the reported polymer type of the microplastic sample from which it originated, as described in the original studies. In total, six distinct polymer types were identified across all samples, with polypropylene (PP) being the most reported, representing 21.74% of the total (n = 10/46 metagenomic read libraries). This was followed by polycaprolactone (PCL) and polyvinyl chloride (PVC), each accounting for 15.22% (n = 7/46) and 13.04% (n = 6/46), respectively. The full distribution of microplastic types across the datasets is shown in **Fig. 3A**. The taxonomic profiles of microbial communities associated with these microplastic types revealed notable differences in phylum-level composition (**Fig. 3B**). Across all microplastic types, the phylum *Pseudomonadota* dominated the communities, particularly on PP, polylactic acid (PLA), and polystyrene (PS), where they constituted most of the detected taxa. In contrast, PCL exhibited a distinct signature characterized by *Desulfobacterota* and *Patescibacteria* relative to the other polymers; however, PCL data were available only from one study. Further taxonomic analysis of the recovered MAGs identified several genera known to harbor pathogenic or opportunistic bacteria associated with both marine organisms and humans **(Table S3)**. Genera such as *Vibrio* (Baker-Austin *et al*. 2018; Marques *et al*. 2022)*, Pseudoalteromonas* (Pujalte *et al*. 2007)*, Tenacibaculum* (Mabrok *et al*. 2022)*, Psychromonas* (Wei *et al*. 2024)*, Flavobacterium* (Loch and Faisal 2015; Zurbuchen *et al*. 2023), and *Shewanella* (Paździor 2016; Yu *et al*. 2022) were detected across multiple microplastic types.

**Fig. 3.**
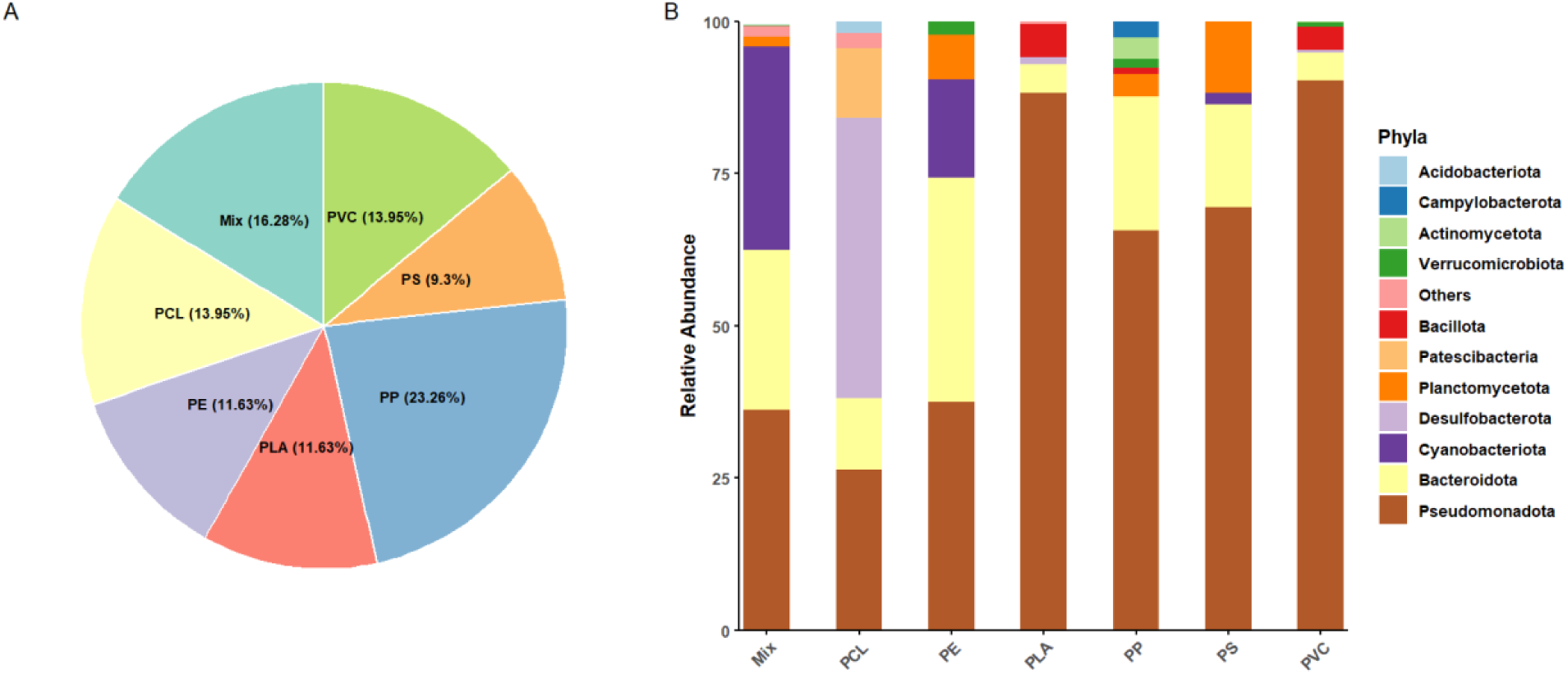
(A) Distribution of microplastic types in the metagenomic dataset. (B) Relative abundance of phylum-level taxonomic composition of microbial communities associated with each MP type based on the metagenomic analysis. These relative abundances represent the proportion of reads mapped to the MAGs assigned to each bacterial phylum within the microbial communities colonizing each microplastic type. Microplastic types were derived from multiple independent studies and samples: PP (3 studies; 20 samples), PE (3 studies; 20 samples), PS (2 studies; 7 samples), PVC (2 studies; 6 samples), PLA (1 study; 3 samples), and PCL (1 study; 3 samples). “Mix” refers to samples in which multiple polymer types were present within the same sequencing library and could not be resolved to a single dominant polymer.

### Distribution of Predicted Plasmid Mobility Across Microplastic Types

The analysis of predicted plasmid mobility revealed varied patterns across the six MP types (**Fig. 4; Table S4)**. A total of 871 replicons were predicted in the plasmid sequences from the geNomad predictions, with varying numbers across the 6 studies **(Table S5).** Overall, conjugative plasmids, which have a complete set of genes required for self-mediated cell-to-cell transfer, were the mostly commonly recovered plasmid type in most microplastic-associated communities, accounting for over half of the plasmid content in PE (63%, n = 206/325), PVC (55%, n = 117/211), and PP (53%, n = 55/103). Mobilizable plasmids, lacking a full conjugation system but that can be transferred in the presence of co-resident conjugative elements, were frequently observed on PS, accounting for 72% (n = 57/79) of the recovered plasmids. PLA-associated communities harbored the highest fraction of non-mobilizable plasmids, which lack mobility and transmission genes (65%, n = 73/113), with notable numbers also recovered from PP (40%, n = 41/103) and PE (36%, n = 117/325).

**Fig. 4.**
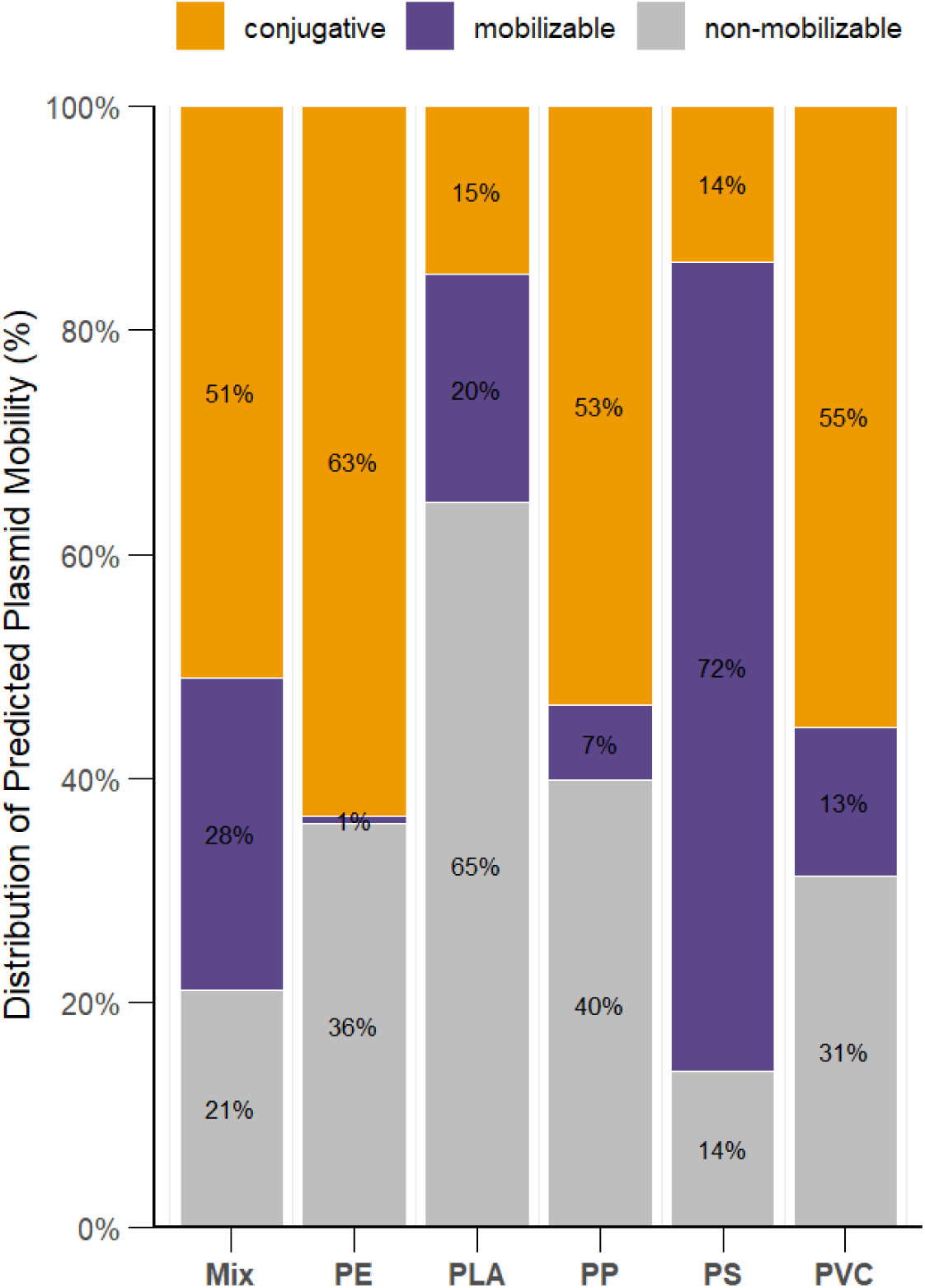
Distribution of predicted plasmid mobility types across different plastic types. Plasmid contigs were identified from MAGs using the NMDC viruses and plasmid workflow, and mobility classification was performed using the Plasmid Database (PLSDB). Each bar represents the relative proportion of predicted plasmids categorized as conjugative (orange), mobilizable (purple), or non-mobilizable (gray) for each plastic type. Percentages are based on the total number of plasmids identified per plastic type.

### Co-occurrence of Antibiotic Resistance Genes and Plasmid Replicons

Analysis of plasmid-associated ARGs revealed a diverse distribution of resistance determinants across different plasmid replicons (**Fig. 5**). Among the detected plasmids, members of the IncF family were the most observed, co-occurring with genes conferring resistance to multiple antibiotic classes such as aminoglycosides and beta-lactams.

**Fig. 5.**
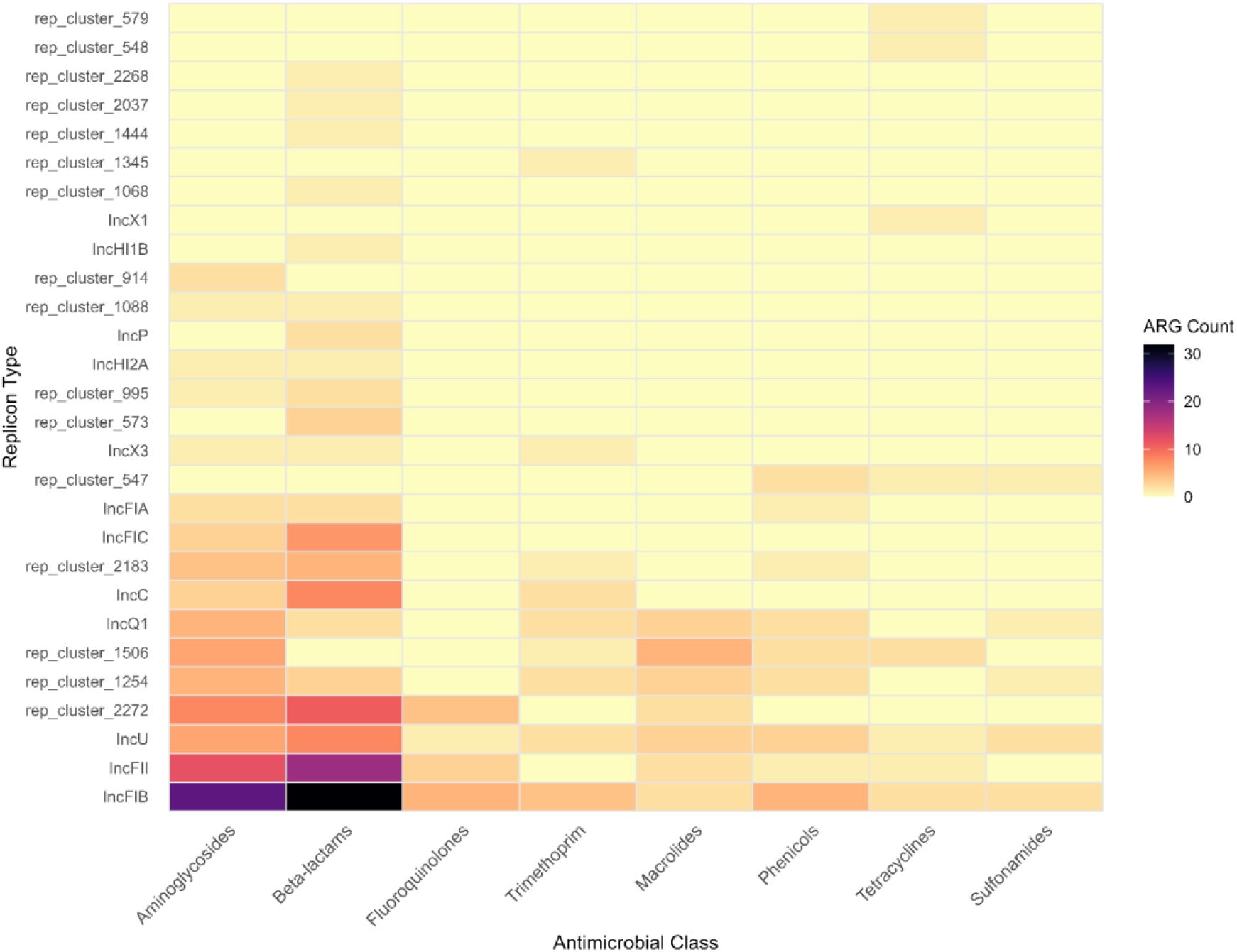
Heatmap showing the number of plasmid replicon types associated with different antimicrobial classes. The rows represent the plasmid replicons identified using PLSDB, while the columns represent the major ARG classes. Color intensity reflects the count of plasmid-ARG associations, with darker shades indicating higher co-occurrence.

### Distribution of ARG Classes Across Plasmid Replicons and Mobility Types

Figure 6 illustrates the connectivity between plasmid mobility categories, the top 25 most observed plasmid replicons, and their associated antimicrobial drug classes.

**Fig. 6.**
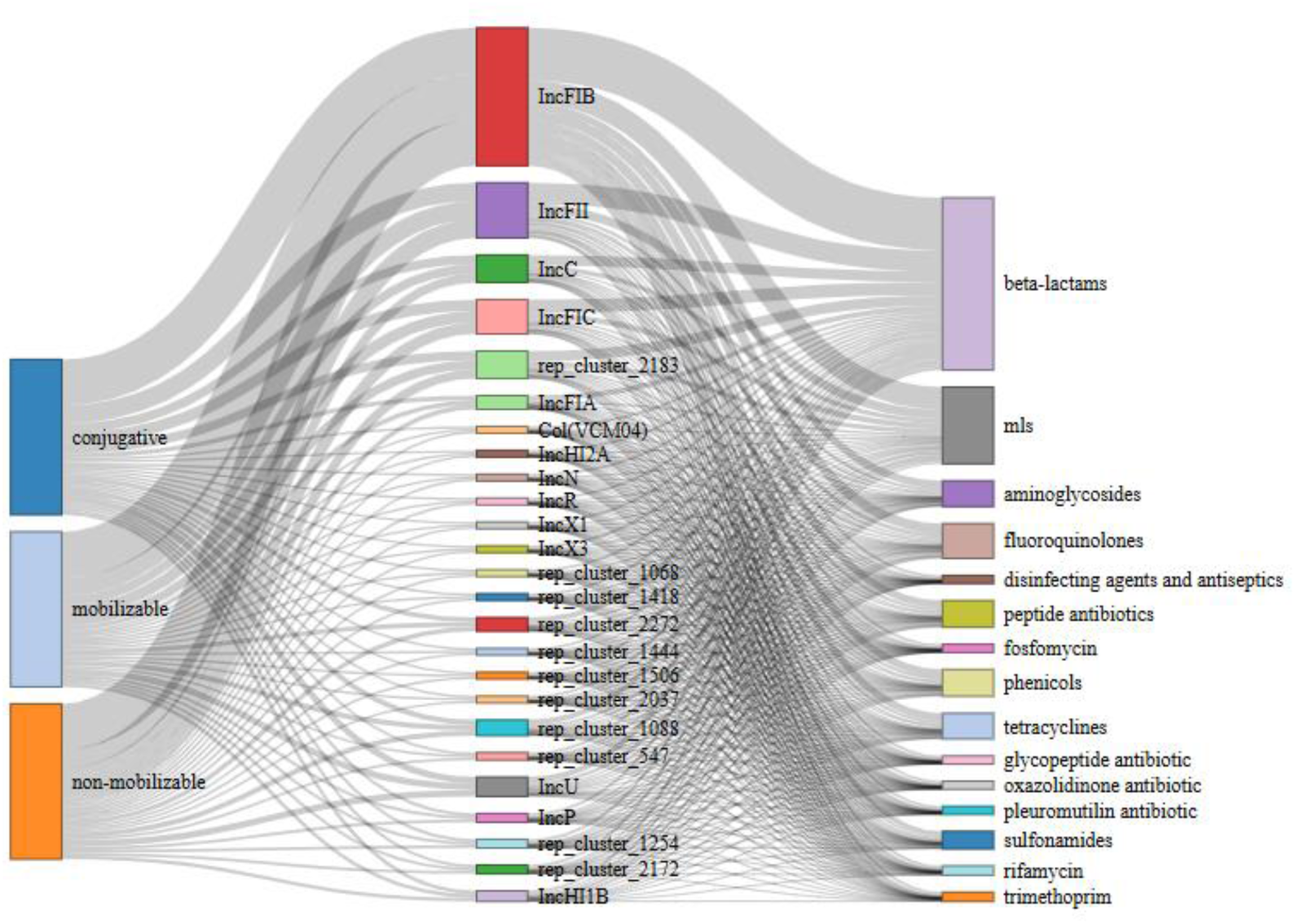
Sankey diagram illustrating the connections between plasmid replicon types, their predicted mobility classifications, and associated ARG class. The diagram links (left) plasmid mobility types, (center) the top 25 most dominant plasmid replicons within the dataset, and (right) ARG classes co-occurring with the plasmids. Mobility groups are categorized as conjugative, mobilizable, or non-mobilizable, while individual replicons are color-coded to emphasize the most abundant plasmid types. The drug classes represented include major clinically relevant groups, such as beta-lactams, MLS (macrolides, lincosamides, streptogramins), aminoglycosides, tetracyclines, peptide antibiotics, and others.

Conjugative plasmids were associated with most ARGs (153 ARGs across 46 plasmids), followed by non-mobilizable (135 ARGs across 26 plasmids) and mobilizable plasmids (15 ARGs across 10 plasmids), reflecting their differing capacities for horizontal gene transfer. Within the replicon layer, the well-characterized incompatibility replicons of the Inc plasmid family (IncFIB, IncFII, IncC, IncFIC, IncFIA) dominated the dataset. In addition, less-characterized replicons, the rep_cluster groups (e.g., rep_cluster_1068, rep_cluster_1418, rep_cluster_2183), were also frequently observed. Across drug classes, beta-lactams, macrolides-lincosamides-streptogramins (MLS), aminoglycosides, fluoroquinolones, and tetracyclines were the most widely represented and associated with the Inc replicon types.

## DISCUSSION

The composition of microplastics analyzed in this study (Fig. 2A) reflects the diverse range of plastic polymers commonly detected in aquatic environments, with PP as the most frequently reported. This is consistent with previous findings that have identified PP, PS, and PE as the most abundant plastic types in aquatic environments (Di and Wang 2018; Eriksen *et al*. 2013; Matsuguma *et al*. 2017) due to their widespread use in consumer packaging, cosmetics, and care products (Fendall and Sewell 2009).

The microbial community composition associated with these polymers (Fig. 2B) shows consistent taxonomic dominance by *Pseudomonadota* across all MP types, albeit with varying relative abundances. This observation aligns with multiple studies reporting that *Pseudomonadota* are among the most frequently recovered phyla from microplastic surfaces (Wang *et al*. 2020; Witsø *et al*. 2025; Wu *et al*. 2019b). A key factor contributing to their dominance is the ability of many *Pseudomonadota* taxa to produce abundant extracellular polymeric substances and surface-adhesive structures, which enhance initial attachment and biofilm maturation on hydrophobic substrates. These traits likely provide a competitive advantage on plastic surfaces, enabling *Pseudomonadota* to establish and persist as core members of plastisphere communities (Tang 2024).

Our data also highlighted substantial relative abundances from *Bacteroidota* and *Cyanobacteriota*. The presence of *Bacteroidota*, known for their role in organic matter degradation, is consistent with prior observations of their enrichment in MP-associated biofilms, particularly in nutrient-rich or anthropogenically impacted waters (Fortin *et al*. 2025; Ventura *et al*. 2024). *Cyanobacteriota*, on the other hand, have been noted in plastisphere studies from both freshwater and marine systems, likely favored by the hydrophobic surfaces and light availability in surface waters (de Oliveira *et al*. 2020; Zhai *et al*. 2023). While our findings confirm many known trends in plastisphere composition, they also reveal subtle differences in taxonomic composition across polymer types, such as *Desulfobacterota* and *Patescibacteria* being more abundant on PCL than the other polymer types. Despite this, uncertainty exists over how strongly polymer type drives microbial differentiation (Bhagwat *et al*. 2021; Jacquin *et al*. 2019; Miao *et al*. 2023). While certain studies report distinct community profiles on specific plastics (McCormick *et al*. 2016; Mughini-Gras *et al*. 2021), others find that communities frequently converge despite variation in polymer properties. This inconsistency likely reflects the interaction between polymer characteristics and environmental processes.

Features such as surface roughness, topography, electrostatic charge, and hydrophobicity strongly influence initial bacterial adhesion and early biofilm development (Rummel *et al*. 2017). Environmental aging processes, including weathering, oxidation, and photo-induced chemical modification, further modulate these properties by altering surface energy, creating micro-textures, and exposing functional groups that enhance microbial colonization (Bao *et al*. 2022; Likpalimor *et al*. 2025). These mechanisms could explain the distinct bacterial assemblage observed on PCL in our study. Unlike conventional, non-degradable plastics, PCL is a biodegradable polyester that undergoes hydrolytic and microbial breakdown. This degradation increases surface roughness, reduces hydrophobicity, and releases low-molecular-weight carbon substrates, collectively creating microhabitats that differ from those found on more inert polymers (Mu *et al*. 2023). Such conditions can favor the proliferation of specialized taxa, including *Desulfobacterota* and *Patescibacteria*, which were not comparably observed on the other plastics.

The potential hazards associated with microplastics can be broadly categorized into three main aspects: (1) the inherent toxicity of the plastic particles themselves, (2) the toxicity of chemical contaminants adsorbed onto microplastic surfaces, and (3) the colonization of MPs by pathogenic microorganisms (Auta *et al*. 2017; Noventa *et al*. 2021). In this study, several taxa identified on microplastics are known marine pathogens, including *Vibrio, Pseudoalteromonas, Tenacibaculum, Psychromonas, Flavobacterium*, and *Shewanella* (Johnson *et al*. 2025; Richards *et al*. 2008; Rodgers *et al*. 2014), and have been implicated in infections affecting fish, shellfish, and other marine organisms (Farto *et al*. 2003; Loch and Faisal 2015), underscoring the ecological and aquaculture-related risks of microplastic-associated microbial communities. The detection of these genera on microplastics does not necessarily indicate active infection risk. Rather, it reflects the ability of microplastics to serve as stable nutrient-enriched microhabitats that support the persistence and proliferation of potentially harmful microbes. Additionally, since the gastrointestinal tract is a primary route of bacterial infection in fish and invertebrates, ingestion of contaminated microplastics could increase exposure risk and facilitate the spread of infections in natural and aquaculture systems (Ghosh 2024). Also, several of these taxa, most notably *Vibrio* and *Shewanella*, can cause opportunistic infections in humans, particularly in immunocompromised individuals. These genera have been implicated in skin and soft tissue infections, including cellulitis and wound-associated infections, as well as infections acquired through seafood consumption or direct contact with marine environments (Di Bartolomeo *et al*. 2025). Thus, their detection on microplastics suggests that plastic-associated biofilms may act as environmental reservoirs that support the persistence of bacteria with dual relevance to marine ecosystem health and human health.

Dense biofilms increase cell-to-cell contact rates, which can enhance genetic exchange, particularly among taxa such as *Vibrio* that readily engage in HGT under favorable conditions (Aminov 2011). In this context, the detection of mobile genetic elements within plastisphere communities strengthens the idea that microplastics may function as microenvironments that facilitate HGT between environmental bacteria and opportunistic pathogens. The plasmid profiling further supports this interpretation: distinct mobility patterns were observed across plastic types, with conjugative plasmids commonly recovered from biofilms on PE, PVC, and PP—polymers that also hosted distinct and taxonomically diverse microbes. Conjugative plasmids encode the machinery required for self-transfer. Thus, their dominance on these surfaces suggests that microplastic-associated communities may possess an enhanced capacity for genetic exchange within mixed-species assemblages. This is ecologically relevant, as conjugative plasmids frequently carry accessory genes linked to stress tolerance, metabolic flexibility, and antimicrobial resistance, traits that can enhance bacterial persistence in heterogeneous and dynamic surface-associated environments such as microplastics.

Accordingly, conjugative plasmids also emerged as the primary conduits linking plasmid replicons to ARG classes across the network (Fig. 6), reflecting both their broad host range and established role as major vectors for the horizontal dissemination of clinically important resistance determinants, including to beta-lactams, MLS antibiotics, and aminoglycosides. Several of the ARGs are of particular concern from a One Health perspective **(Table S6)**. These include extended-spectrum and carbapenemase genes such as *bla*_CTX-M-3_, *bla*_TEM-1_, *bla*_OXA-1_, and notably *bla*_KPC-2_, a globally disseminated carbapenem resistance determinant that severely limits treatment options for Gram-negative infections (Lee *et al*. 2016). The detection of *bla*_KPC-2_ on microplastic-associated plasmids is especially noteworthy given its strong association with healthcare-associated outbreaks and multidrug-resistant Enterobacterales (Kerdsin *et al*. 2019).

Resistance determinants targeting fluoroquinolones and aminoglycosides were also prevalent, including *qnrVC4*, *qnrVC5*, *aac(6)-Ib-cr*, *aadA2*, and *adeF*, which collectively confer reduced susceptibility to first-line antibiotics commonly used in both clinical and veterinary medicine. In addition, the presence of *dfrA* variants (*dfrA19, dfrA31*), *sul1*, and *sul2* highlights resistance to folate pathway inhibitors, drugs that are widely used and frequently detected in aquatic environments (Chaturvedi *et al*. 2021). The co-occurrence of clinically relevant ARGs with conjugative plasmids on microplastic surfaces suggests that the plastisphere may act as an environmental reservoir in which resistance determinants of direct human health relevance are concentrated within highly mobile genetic platforms.

Within this framework, the incompatibility family replicons, particularly IncFIB and IncFII, emerged as prominent hubs connecting multiple ARG classes across mobility categories. These replicons are well-documented in both environmental and clinical contexts, especially among Enterobacterales, where they are frequently associated with multidrug-resistant phenotypes (Carattoli 2009, 2013; Rozwandowicz *et al*. 2018). Their strong representation in the plastisphere communities suggests that microplastic-associated resistomes can be shaped by plasmids commonly implicated in both environmental and clinically relevant resistance dissemination, consistent with a One Health perspective in which resistance genes circulate across different ecological compartments. Notably, IncFIB and IncFII plasmids were the replicon types most frequently associated with ARGs in this study, with strong associations to beta-lactam and aminoglycoside resistance genes (Fig. 5). IncF plasmids are known for their large, flexible accessory regions, which facilitate the accumulation and maintenance of multiple resistance determinants (Osborn *et al*. 2000). Hence, their presence suggests that the plastisphere may provide favorable conditions for the persistence of such plasmids, potentially due to prolonged surface attachment and biofilm stability. In parallel, several less-characterized rep_cluster replicons (e.g., rep_cluster_1068, rep_cluster_1418, rep_cluster_2183) were frequently detected and associated with multiple ARG classes, including sulfonamides and tetracyclines. Although the mobility potential and host range of these replicons remain poorly resolved, their repeated occurrence suggests that microplastic-associated biofilms may harbor a diverse pool of environmentally adapted plasmid backbones that can maintain resistance genes. Rather than acting as dominant vectors of horizontal transfer, these less-characterized plasmids may function as stable reservoirs of ARGs within plastisphere communities, highlighting the role of microplastics as sites of resistome accumulation and persistence.

## CONCLUSIONS

This comparative multi-study, metagenomic analysis demonstrates that microplastic surfaces consistently support distinct microbial assemblages and plasmid-associated resistomes across diverse polymer types. Despite originating from independent studies and environmental contexts, plastisphere communities shared common taxonomic and functional features, including the dominance of surface-adapted bacterial phyla and a high prevalence of plasmid-encoded ARGs. These findings reinforce the view of microplastics as stable, long-lived microbial habitats that differ from surrounding environments in both community structure and functional potential.

A key contribution of this study is the characterization of plasmids underlying plastisphere resistomes. Conjugative plasmids formed the primary links between plasmid replicons and clinically relevant ARG classes, highlighting their central role as genetic platforms that facilitate resistance dissemination within surface-associated microbial communities. The strong representation of Inc family replicons, particularly IncFIB and IncFII, mirrors well-established patterns in both environmental and clinical microbiology and underscores the relevance of plastisphere-associated plasmids to broader resistance gene circulation. At the same time, the frequent detection of less-characterized rep_cluster replicons suggests that microplastic-associated biofilms also harbor diverse, environmentally adapted plasmid backbones that may function as reservoirs for resistance determinants, even in the absence of confirmed mobility.

Importantly, several ARGs detected in this study, including extended-spectrum and carbapenem-associated β-lactamases, and fluoroquinolone resistance genes, are of recognized public-health concern. The co-occurrence of these genes with mobile plasmid frameworks on microplastic surfaces highlights the potential for microplastics to concentrate resistance determinants within diverse environmental niches.

## DATA AVAILABILITY STATEMENT

The data supporting the findings of this study are available within the article and/or the Supplementary Tables.

## ACKNOWLEDGEMENTS

This work was supported by a NYS Center of Excellence in Healthy Water Solutions Summer Fellowship to CT. This material is based upon work supported by the U.S. Geological Survey under Grant/Cooperative Agreement No. G21AP10626-01.

## AUTHOR CONTRIBUTIONS STATEMENT

Conceptualization: IJO, JLG; Methodology: IJO, CT, JLG; Validation: IJO, JLG; Formal Analysis: IJO; Investigation: IJO, CT; Data Curation: IJO, CT; Writing – Original Draft: IJO; Writing – Review & Editing: IJO, CT, JLG; Visualization: IJO; Supervision: JLG; Funding Acquisition: JLG

